# Reward timing matters in motor learning

**DOI:** 10.1101/2021.10.15.464563

**Authors:** Pierre Vassiliadis, Aegryan Lete, Julie Duque, Gerard Derosiere

**Affiliations:** Institute of Neuroscience, Université Catholique de Louvain, 1200, Brussels, Belgium; Defitech Chair for Clinical Neuroengineering, Center for Neuroprosthetics (CNP) and Brain Mind Institute (BMI), Swiss Federal Institute of Technology (EPFL), 1202, Geneva, Switzerland

**Keywords:** Motor learning, motor skill learning, memory consolidation, reward, motivation, reinforcement learning, timing

## Abstract

Reward can improve motor learning and the consolidation of motor memories. Identifying the features of reward feedback that are critical for motor learning is a necessary step for successful integration into rehabilitation programs. One central feature of reward feedback that may affect motor learning is its timing – that is, the delay after which reward is delivered following movement execution. In fact, research on associative learning has shown that short and long reward delays (*e.g*., 1 and 6 s following action execution) activate preferentially the striatum and the hippocampus, respectively, which both contribute with varying degrees to motor learning. Given the distinct functional role of these two areas, we hypothesized that reward timing could modulate how people learn and consolidate a new motor skill. In sixty healthy participants, we found that delaying reward delivery by a few seconds influenced motor learning dynamics. Indeed, training with a short reward delay (*i.e*., 1 s) induced slow, yet continuous gains in performance, while a long reward delay (*i.e*., 6 s) led to initially high learning rates that were followed by an early plateau in the learning curve and a lower endpoint performance. Moreover, participants who successfully learned the skill with a short reward delay displayed overnight consolidation, while those who trained with a long reward delay exhibited an impairment in the consolidation of the motor memory. Overall, our data show that reward timing affects motor learning, potentially by modulating the engagement of different learning processes, a finding that could be exploited in future rehabilitation programs.

## 1. Introduction

When delivered following well-executed movements, reward can boost motor learning (Chen et al., 2017; Dhawale et al., 2017; Galea et al., 2015; Vassiliadis et al., 2021) and the consolidation of motor memories (Abe et al., 2011). This observation has raised hope for rehabilitation, where reward is regarded as a promising means to magnify the positive effects of practice on motor control (Quattrocchi et al., 2017; Therrien et al., 2020, 2016; Vassiliadis et al., 2019; Vassiliadis and Derosiere, 2020). Yet, this branch of research is only burgeoning, and a current challenge in the field is to identify the features of reward feedback that may be critical for motor learning.

Recent studies have started to tackle this issue, showing that the magnitude (Vassiliadis et al., 2021), the valence (Galea et al., 2015) and the stochasticity (Dayan et al., 2014) of reward feedback bear all a decisive impact on motor learning. Another key feature of reward feedback that may directly affect motor learning is its timing – that is, the delay after which reward is delivered following movement execution. As such, previous studies have shown that reward prediction error signals, which are key for reward-based learning, are not only modulated by the value of the reward but also depend on the timing at which it is delivered (Fiorillo et al., 2008; Klein-Flügge et al., 2011; Kobayashi and Schultz, 2008). Moreover, converging lines of evidence from neuroimaging and electroencephalographic studies indicate that different brain structures – known to be involved with varying degrees in motor learning – exhibit activity changes in response to reward feedback depending on its timing. Indeed, in associative learning tasks, short reward delays (*e*.*g*., provided 1 s following action execution) activate a fronto-striatal network, while long reward delays (*e*.*g*., 6 s following execution) evoke changes in the activity of the hippocampus primarily (Foerde and Shohamy, 2011; Peterburs et al., 2016). Further, Parkinson’s disease and ADHD patients, both known to exhibit striatal dysfunction (Mehler-Wex et al., 2006), are impaired in learning action-outcome associations based on short reward delays (Foerde and Shohamy, 2011; Gabay et al., 2018; Weismüller et al., 2018), while amnesic patients with damage to the hippocampus are unable to learn associations with long reward delays (Foerde et al., 2013). Altogether, these findings indicate that the processing of reward preferentially engages striatum-or hippocampus-centred networks depending on the timing at which it is delivered.

The striatum and the hippocampus show varying contributions during motor learning and consolidation (Doyon and Benali, 2005; Fernández-Seara et al., 2009; Krakauer et al., 2019; Schendan et al., 2003), which are thought to underlie the operation of distinct learning processes (Albouy et al., 2013, 2008). Hence, it is sensible to assume that reward may boost different motor learning processes – potentially relying on the striatum or the hippocampus – depending on the timing at which it is delivered. Notably, previous studies on reward-based motor learning have only exploited short reward delays, preventing one to test this hypothesis directly. Here, we tested this idea by evaluating the performance of sixty healthy participants in a skill learning task (Vassiliadis et al., 2021), where reward was delivered either at a short or at a long delay following movement execution. We found that delaying reward delivery by a few seconds influenced the dynamics of learning. Indeed, training with a short reward delay induced slow, yet continuous gains in performance, while a long reward delay led to initially high learning rates that were followed by an early plateau in the learning curve and a lower endpoint performance. Moreover, participants who successfully learned the skill with a short reward delay displayed overnight consolidation, while those who trained with a long reward delay exhibited an impairment in the consolidation of the motor memory. Altogether, the present results provide evidence that reward timing can strongly influence motor learning, a finding that could be exploited in future rehabilitation protocols.

## 2. Results

Sixty healthy participants practiced a pinch-grip force task over two consecutive days. Participants were required to hold a pinch grip transducer in their right hand and to squeeze it as quickly as possible in order to move a cursor displayed on a computer screen in front of them, from an initial position to a fixed target (**Figure 1A**; (Vassiliadis et al., 2021)). The force required to reach the target (Target_Force_) corresponded to 10 % of the individual maximum voluntary contraction (MVC). In most of the trials (90 %), participants practiced the task with very limited sensory feedback: the cursor disappeared when the generated force reached half of the Target_Force_ (see STAR Methods for more details on the task). To learn the task, subjects were provided with six Training blocks (T1 to T6; 40 trials each; *i*.*e*., total of 240 training trials; **Figure 1B**) in which they received reinforcement feedback (*i*.*e*., indicating Success or Failure) associated to a monetary reward. Success on the task was determined based the Error, defined as the absolute force difference between the Target_Force_ and the exerted force (Abe et al., 2011; Steel et al., 2016).

**Figure 1.**
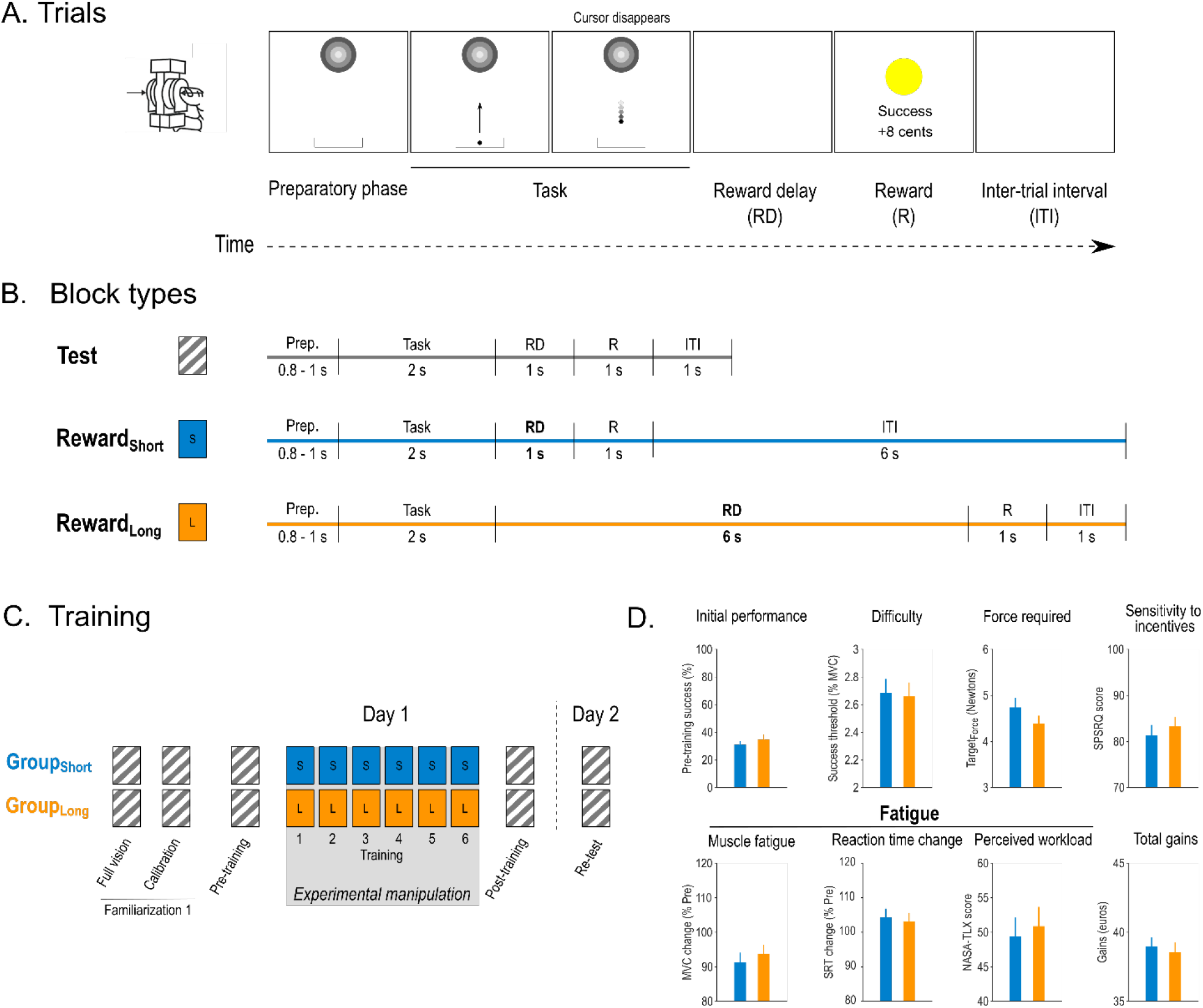
Motor skill learning task. **A. Time course of a trial in the motor skill learning task**. Each trial started with the appearance of a sidebar and a target. After a variable preparatory period (0.8-1s), a cursor appeared in the sidebar, playing the role of a “Go” signal. At this moment, participants were required to pinch the force transducer to bring the cursor into the target as quickly as possible and maintain it there until the end of the task (2 s). Notably, on most trials, the cursor disappeared halfway towards the target (as displayed here). Then, after a delay, a reward (R) appeared consisting in a reinforcement feedback and a monetary reward (a successful trial is shown here). Trials ended with an inter-trial interval (ITI). **B. Durations in the different block types**. Reward delays (RD) and ITIs were manipulated. Test blocks included a short reward delay (1 s), a short ITI (1 s) and no monetary reward (*i*.*e*., only reinforcement). Reward_Short_ and Reward_Long_ blocks included monetary rewards and were performed with a short (1 s) and long (6 s) reward delay, respectively. The total duration of the trials was kept constant between Reward_Short_ and Reward_Long_ by varying the ITI. **C. Training procedure**. On Day 1, all participants performed two familiarization blocks in a Test blocks condition. The first one involved full vision of the cursor while the second one provided only partial vision and served to calibrate the difficulty of the task on an individual basis (See STAR Methods). Then, Pre- and Post-training Test blocks assessments were separated by 6 blocks of training in the condition corresponding to each individual group (Reward_Short_ for Group_Short_, Reward_Long_ for Group_Long_). Day 2 consisted in a short re-familiarization (5 trials with full vision, not represented) followed by a Re-test assessment (1 Test block). **D. Control analyses**. Group_Short_ and Group_Long_ were comparable for a variety of factors including initial performance, task difficulty, required force to reach the target, sensitivity to reward and punishment (as assessed by the SPSRQ questionnaire), muscular and cognitive fatigue and final monetary gains (see also Table 1).

In different groups of participants, we varied the delay between the end of the movement period and the delivery of the reward during the Training blocks. As such, Group_Short_ subjects trained with a short reward delay (*i*.*e*., 1 s) while participants of the Group_Long_ performed the task with a long reward delay (*i*.*e*., 6 s). The total duration of the trials was kept constant by modulating the inter-trial interval (ITI; 6 s in Group_Short_ and 1 s in Group_Long_). Before, immediately and 24 hours after training, all participants performed Test blocks with no reward, a short reward delay (1 s) and a short ITI (1 s). Notably, the groups were comparable for a variety of features including Pre-training success rates, difficulty of the task, force required, sensitivity to reward and punishment, fatigue and final monetary gains (**Figure 1C, Table 1**). Altogether, this design allowed us to investigate the specific effect of reward timing on motor learning and consolidation.

**Table 1.**
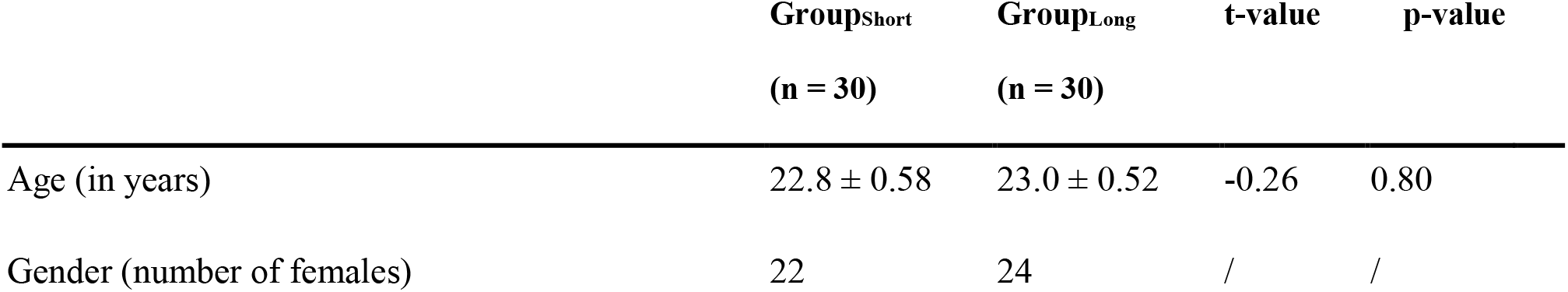
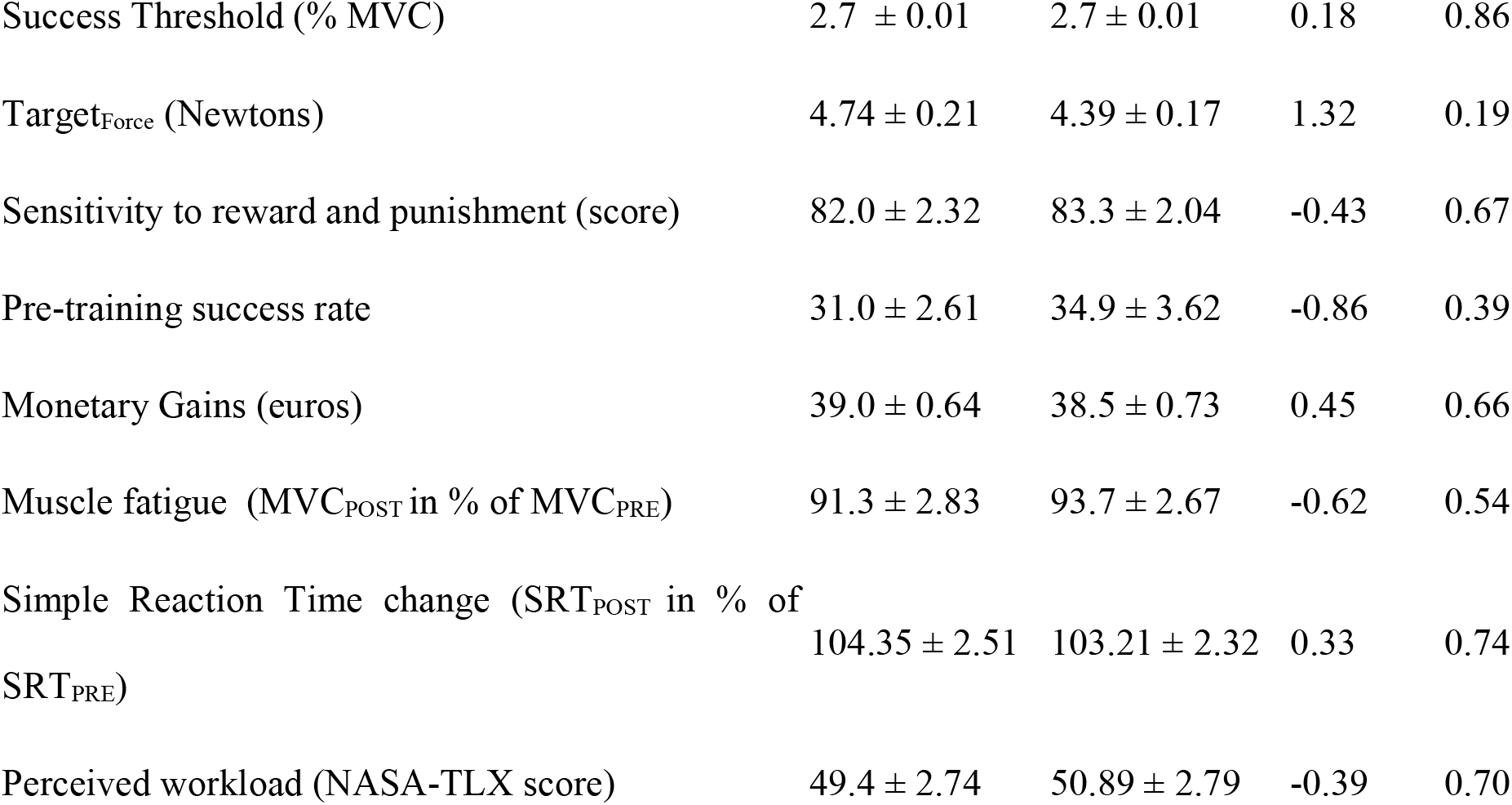
Group features, initial performance and fatigue in the three experimental groups (mean ± SE). The 2 last columns provide the results of independent samples t-tests.

|  | Group<sub>Short</sub><br>(n = 30) | Group<sub>Long</sub><br>(n = 30) | t-value | p-value |
| --- | --- | --- | --- | --- |
| Age (in years) | 22.8 ± 0.58 | 23.0 ± 0.52 | -0.26 | 0.80 |
| Gender (number of females) | 22 | 24 | / | / |
| Success Threshold (% MVC) | $2.7 \pm 0.01$ | $2.7 \pm 0.01$ | 0.18 | 0.86 |
| Target<sub>Force</sub> (Newtons) | $4.74 \pm 0.21$ | $4.39 \pm 0.17$ | 1.32 | 0.19 |
| Sensitivity to reward and punishment (score) | $82.0 \pm 2.32$ | $83.3 \pm 2.04$ | -0.43 | 0.67 |
| Pre-training success rate | $31.0 \pm 2.61$ | $34.9 \pm 3.62$ | -0.86 | 0.39 |
| Monetary Gains (euros) | $39.0 \pm 0.64$ | $38.5 \pm 0.73$ | 0.45 | 0.66 |
| Muscle fatigue (MVC<sub>POST</sub> in % of MVC<sub>PRE</sub>) | $91.3 \pm 2.83$ | $93.7 \pm 2.67$ | -0.62 | 0.54 |
| Simple Reaction Time change (SRT<sub>POST</sub> in % of<br>SRT<sub>PRE</sub>) | $104.35 \pm 2.51$ | $103.21 \pm 2.32$ | 0.33 | 0.74 |
| Perceived workload (NASA-TLX score) | $49.4 \pm 2.74$ | $50.89 \pm 2.79$ | -0.39 | 0.70 |

### Training with long reward delays induces an early plateau in learning and a reduction of endpoint performance

As a first step, we evaluated performance on the task by computing the average success rate per Training_Block_ (T1 to T6). To compare the learning process between the groups, we performed a two-way ANOVA, with TRAINING_BLOCK_ and GROUP_TYPE_ as within- and between-subject factors, respectively. Overall, participants of both groups significantly improved their success rates over training (main effect of TRAINING_BLOCK_: Greenhouse-Geisser [GG]-corrected F_(3.56, 206.77)_ = 6.01; p < 0.001, partial η^2^ = 0.094; **Figure 2A**). Most importantly, the improvement in success rate over the blocks depended on the Group_Type_, as revealed by a significant TRAINING_BLOCK_ x GROUP_TYPE_ interaction (GG-corrected F_(3.56, 206.77)_ = 2.69; p = 0.038, partial η^2^ = 0.044; **Figure 2A**). As such, in Group_Short_, performance only started to improve significantly at T4 relative to T1 (T4 vs. T1: p = 0.044; T5 vs T1: p = 0.030; T6 vs T1: p < 0.001) and kept improving in late Training_Blocks_ (T6 vs T3: p < 0.001; T6 vs T4: p = 0.034; T6 vs T5: p = 0.051). In contrast, Group_Long_ improved early on (*i*.*e*., T2 vs T1: p < 0.001) but then stopped improving quickly during training, with the success rate at T2 being not different from the success rate at any of the following Training_Blocks_ (all p > 0.37). Interestingly, post-hoc tests further revealed that endpoint performance (*i*.*e*., success rate at T6) was significantly lower in Group_Long_ than in Group_Short_ (p = 0.047; **Figure 2B**). Conversely, success rates at all other Training_Blocks_ were comparable between the two groups (all p > 0.22; **Figure 2A**). Altogether, these findings suggest that reward timing can influence the dynamics of motor learning. More specifically, it appears that training with long reward delays is associated with an early plateau in learning and a related reduction of endpoint performance.

**Figure 2.**
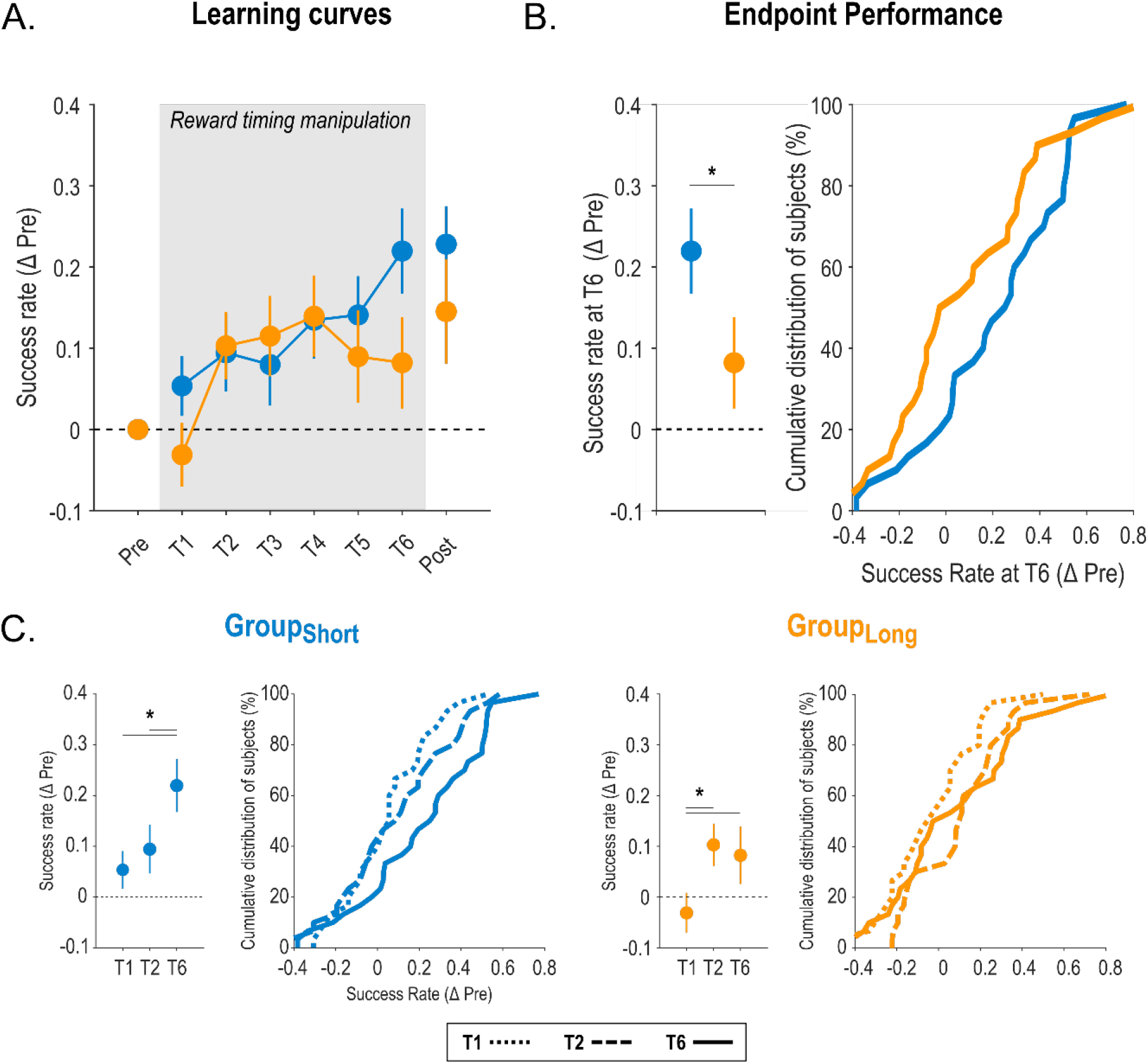
Effect of reward timing on motor skill learning. **A. Learning curves.** Proportion of successful trials (expressed as a difference with the individual Pre-training success rate) is represented across practice for the two experimental groups (blue: Group_Short_, orange: Group_Long_). The grey shaded area highlights the blocks concerned by the reward timing manipulation. The remaining blocks were Test blocks. **B. Endpoint performance**. Average success rates in the end of the training period (i.e., measured at T6) in the two groups (left panel) and the corresponding cumulative distributions of the data (right panel). **C. Learning dynamics**. Average success rates observed at T1 (dotted line), T2 (dashed line) and T6 (solid line) and the corresponding cumulative distributions for Group_Short_ (left) and Group_Long_ (right panel). Performance improved very early in Group_Long_ (i.e., between T1 and T2) and then plateaus while it improved later in Group_Short_. Note that performance at T1 was not different between the groups (all p > 0.21). *: significant difference (p<0.05). Data are represented as mean ± SE.

An important aspect of our experimental design is that we increased the duration of the ITI in Group_Short_ relative to Group_Long_ (6 s and 1 s, respectively; **Figure 1B**), in order to match the total duration of the trials in both groups despite differences in reward timing. To verify that such manipulation did not impact learning in our task, we added in the analysis another group of participants, who trained with a short reward delay (0.5 s) and an intermediate ITI (3 s; Group_Short-PastStudy_, n = 30; from (Vassiliadis et al., 2021)). We reasoned that, if differences in learning dynamics were truly driven by differences in reward timing but not by differences in ITI duration, learning in Group_Short-PastStudy_ should be similar than in Group_Short_, and therefore different than in Group_Long_. Consistently, post-hoc tests run on the TRAINING_BLOCK_ x GROUP_TYPE_ interaction (GG-corrected F_(3.78, 328.92)_ = 2.84; p = 0.0020, partial η^2^ = 0.061) showed (1) no difference at any Training_Blocks_ between Group_Short-PastStudy_ and Group_Short_ (all p > 0.23) and (2) a trend for a difference in endpoint performance when comparing Group_Short-PastStudy_ and Group_Long_ (*i*.*e*., p = 0.049 and 0.053 at T5 and T6, respectively), similarly as when comparing Group_Short_ and Group_Long_ (**Supplementary Figure 1**). Hence, these results confirm that a change of a few seconds in ITI duration does not impact motor learning, as previously reported (McDougle et al., 2015), implying that the differences in learning dynamics observed in groups exposed to short and long reward delays did not result from differences in ITI duration but rather from actual differences in reward timing.

To evaluate total learning we computed success rates at Post-training, which was performed in a Test block setting in both groups. Overall, success rates increased by 22.8 ± 4.69 % in Group_Short_, and 14.5 ± 6.42 % in Group_Long_ with respect to Pre-training. Interestingly, success rates at Post-training were significantly different from 0 in Group_Short_ despite Bonferroni correction (cutoff for significance: p = 0.025; t_(29)_ = 4.87, p < 0.001), indicative of a significant improvement in performance with respect to Pre-training. In contrast, success rates at Post-training were not significantly different from 0 in Group_Long_ after Bonferroni correction (t_(29)_ = 2.26, p = 0.031). However, a t-test on these data did not show any significant difference between the Group_TYPES_ (t_(58)_ = 1.05; p = 0.30). Hence, reward timing only induced a subtle change in total learning that did not reach significance when comparing directly the groups.

Results of the first analysis suggest that reward timing influenced the dynamics of the learning process with long reward delays being associated with a fast improvement of performance quickly followed by a plateau. To confirm this, we aimed at directly quantifying the learning rates for early and late phases of training (Training_Early_ and Training_Late_, corresponding to T1-T3 and T4-T6, respectively). To do so, for each subject and each training phase, we split the data into 12 bins of 10 trials, computed the average success rate for each bin and then performed linear regressions over the bins for each Training_Phase_ (*i*.*e*., 12 bins per Training_Phase_; **Figure 3A**). The slopes of the linear fits (*i*.*e*., expressed in ΔSuccess rate per bin of trials) were exploited as a proxy of the learning rate at a given Training_Phase_ (**Figure 3B, C**). A two-way ANOVA did not show any main effect of the factor TRAINING_PHASE_ (F_(1, 58)_ = 2.80; p = 0.10, partial η^2^ = 0.046), nor any effect of GROUP_TYPE_ on the learning rates (F_(1, 58)_ = 0.14; p = 0.71, partial η^2^ = 0.0023). Interestingly though, we found a strong TRAINING_PHASE_ x GROUP_TYPE_ interaction on these data (F_(1, 58)_ = 9.13; p = 0.0037, partial η^2^ = 0.14; **Figure 3A, B, C**). This interaction was driven by the fact that learning rates were comparable across training phases in Group_Short_ (p = 0.34), while they were significantly lower at Training_Late_ than at Training_Early_ in Group_Long_ (p = 0.0016; **Figure 3B, D, E**). Consistently, learning rates were higher in Group_Short_ than in Group_Long_ at Training_Late_ (p = 0.017). This tended to be the opposite at Training_Early_ (p = 0.061). Notably, intercepts of the linear fits were not significantly different between the groups, neither at Training_Early_ (t_(58)_ = 1.07, p = 0.29), nor at Training_Late_ (t_(58)_ = -1.33, p = 0.19), indicating that between-group differences in learning rates could not be accounted for by differences in initial performance. Hence, this analysis confirms that training with short reward delay induces slow, yet continuous gains in performance, while long reward timings engender large initial learning rates that then drop significantly, indicative of a plateau in learning. Overall, the data show that reward timing impacts learning dynamics during a motor training.

**Figure 3.**
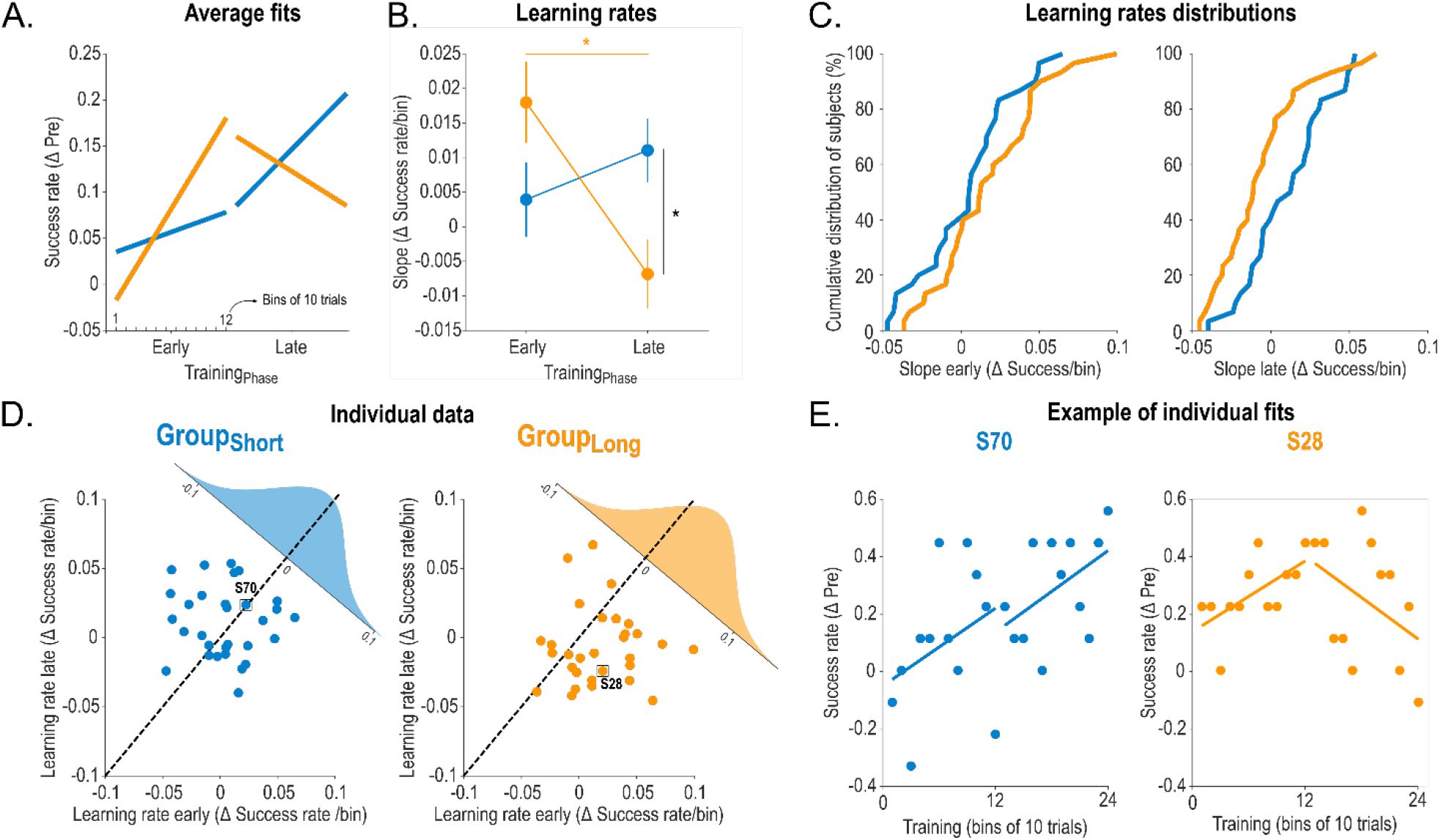
Effect of reward timing on learning rates. **A. Average fits**. Average fits for Group_Short_ (blue) and Group_Long_ (orange) obtained through linear regression on each subject’s binned success data (expressed as a difference from Pre-Training) at early and late phases of training. **B. Learning rates**. The slope of the individual fits – expressed as a delta of success rate per bin on the y-axis – was exploited as a proxy of the learning rate for early and late phases of training. Note the significant TRAINING_BLOCK_ x GROUP_TYPE_ interaction which was driven by the fact that while learning rates were stable in Group_Short_, they were initially high in Group_Long_ and dropped significantly at Training_Late_ (orange star). Moreover, learning rates at Training_Late_ were significantly lower in Group_Long_ than in Group_Short_ (black star). **C. Learning rates distributions**. Cumulative distributions of the group data for early (left panel) and late (right panel) learning phases. **D. Individual learning rates**. Scatter plot representing each subject’s learning rate for early (x-axis) vs late (y-axis) Training_Phases_. The group distributions of the change in learning rates (Training_Early_ – Training_Late_) are also represented (upper right corner of each plot). Note the shift in Group_Long_ below the identity line reflecting the higher learning rates at Training_Early_ than at Training_Late._ **E. Example of individual fits**. Binned success rates and the corresponding linear fits are represented for an exemplar subject of Group_Short_ (left) and Group_Long_ (right). *: significant difference (p<0.05). Data are represented as mean ± SE.

**Figure 4.**
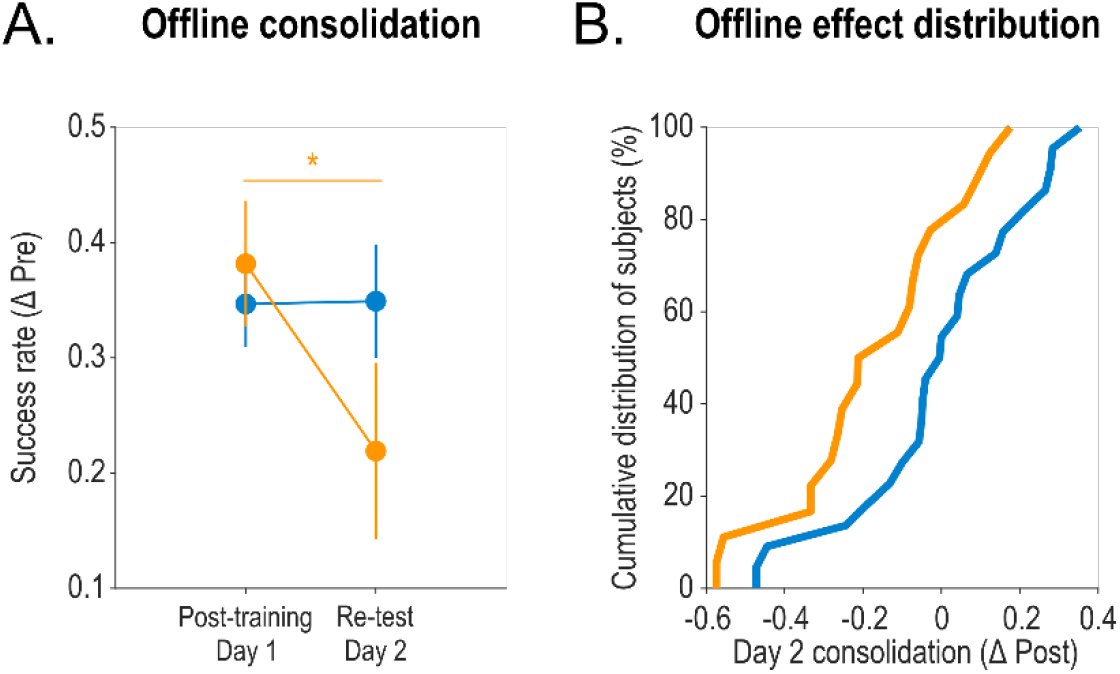
Effect of reward timing on overnight consolidation of the motor skill. **A. Offline consolidation of the motor skill**. Proportion of successful trials (expressed as a difference with the individual Pre-training success rate) at Post-training (Day 1) and Re-test (Day 2). Both assessments were Test blocks. This analysis only considered participants who demonstrated skill learning on Day 1 (n = 22 and 18 in Group_Short_ and Group_Long_, respectively). Notably, a significant TEST_PHASE_ x GROUP_TYPE_ interaction showed that while Group_Short_ participants performed comparably well in Post-training and Re-test, demonstrating offline consolidation of the skill, this process was altered in Group_Long_. **B. Offline effect distribution**. Cumulative distributions of the offline effect (Success rate in Day 2 – Success rate in Post-training) for each Group_Type_

### Training with long reward delay impairs overnight skill consolidation

As a last step, we investigated the impact of the reward timing experienced during training on Day 1 on overnight consolidation of the skill (*i*.*e*., on Day 2). Because this analysis was focused on the consolidation of a learned motor skill, we only included participants who demonstrated learning on Day 1 (n = 22 and 18 in Group_Short_ and Group_Long_, respectively). To evaluate consolidation, we performed a two-way ANOVA on the normalized success rates obtained at Post-training of Day 1 and at Re-test of Day 2 (*i*.*e*., both performed in a Test block setting) with TEST_BLOCK_ and GROUP_TYPE_ as within and between-subjects factors, respectively. The ANOVA showed a significant effect of TEST_BLOCK_ (F_(1, 38)_ = 5.44; p = 0.025, partial η^2^ = 0.13): success rates were on average smaller in the Re-test block on Day 2 than in the Post-training block on Day 1. However, this effect was mostly driven by the Group_Long_, in which performance deteriorated overnight. As such, the ANOVA also revealed a TEST_PHASE_ x GROUP_TYPE_ interaction (F_(1, 38)_ = 5.77; p = 0.021, partial η^2^ = 0.13). While performance was comparable between learners of the two groups at Post-training on Day 1 (p = 0.65), success rates were significantly reduced on Day 2 relative to Day 1 in Group_Long_ (p = 0.0028), but remained stable from one day to another in Group_Short_ (p = 0.95). This indicates that delaying rewards on Day 1 impaired consolidation of the motor skill on Day 2. Overall, our results support the view that short or long reward delays support qualitatively different motor learning processes during training, leading to different consolidation of the skill.

## 3. Discussion

Previous studies have shown that reward timing can influence the response of brain structures involved in reward processing during associative learning (Fiorillo et al., 2008; Foerde et al., 2013; Foerde and Shohamy, 2011; Klein-Flügge et al., 2011; Kobayashi and Schultz, 2008). Inspired by these neurophysiological findings, we asked whether reward timing can also influence how people learn and consolidate a new motor skill. We found that delaying reward delivery by a few seconds influences motor learning dynamics: training with a short reward delay induced slow, yet continuous gains in performance, while a long reward delay allowed high initial learning rates that were followed by an early plateau in the learning curve and a lower endpoint performance. Moreover, among participants who successfully learned the skill, those who trained with a short reward delay displayed overnight consolidation, while those who learned the task with a long reward delay exhibited an impairment in the consolidation of the motor memory. Overall, our findings show that reward timing can influence how the brain learns and consolidates new motor skills.

The differences in learning dynamics observed in subjects trained with short and long reward delays may indicate that reward boosted processes presenting different temporal dynamics. As such, a prevalent view in the field is that motor learning entails the operation of distinct processes, with either slow (*i*.*e*., developing over a few trials) or fast (*i*.*e*., developing over tens/hundreds of trials) temporal dynamics (Smith et al., 2006). The slow process is characterized by both a low learning rate and a sluggish forgetting of the acquired behaviour and is thought to reflect implicit learning (McDougle et al., 2015; Trewartha et al., 2014). In contrast, the fast process entails both a high learning rate and a quick forgetting of the new behavior and supports the explicit learning of new motor behaviors (McDougle et al., 2015; Trewartha et al., 2014). The nature of our task did not allow us to evaluate the relationship between reward timing and the relative contribution of implicit and explicit learning. Still, people who trained with a short reward delay exhibited learning dynamics that presented a low learning rate and a clear overnight consolidation – reminiscent of the slow process, while those who trained with a long reward delay exhibited a high initial learning rate and an overnight forgetting of the motor memory – evocative of the fast process. Based on these results, one may suggest that short reward delays preferentially facilitate the slow (implicit) process, while long reward delays may favour the fast (explicit) learning process, accentuating their respective contribution to subjects’ improvements, an interesting issue for future investigation.

Interestingly, neurophysiological data also support the latter interpretation. Indeed, the striatum – which is preferentially recruited by short reward delays (Foerde and Shohamy, 2011) – exhibits slow changes in activity over the course of motor learning (Albouy et al., 2013, 2008) and, relatedly, was shown to be central for slow, implicit learning (Albouy et al., 2013; Hayes et al., 2015; Siegert et al., 2006). Conversely, the hippocampus – which is mostly recruited by long reward delays – usually exhibits a fast increase in activity in the early phase of learning that wanes later on (Albouy et al., 2013, 2008). In this line, the hippocampus was evidenced to be central for fast, explicit learning (Döhring et al., 2017). Altogether, these elements suggest that the different learning dynamics observed in individuals training with short and long reward delays could result from the fact that these delays recruit the striatum and the hippocampus, respectively, shaping thus their activity and boosting preferentially the slow and fast processes in which those structures are mostly involved. Further neurophysiological work is required to test this hypothesis directly.

The impairment of motor consolidation observed in subjects who trained with a long reward delay also suggests that the strength of learning was altered in these participants. As such, efficient reward-based motor learning relies on the mapping between somatosensory sensations (*e*.*g*., elicited by the generated force in the present task) and the associated reward (Bernardi et al., 2015; Sidarta et al., 2016; Vassiliadis et al., 2021), and somatosensory working memory is known to decay quickly following movement execution, after only a few seconds (Harris et al., 2001; Sidarta et al., 2018). Hence, it is possible that delaying reward delivery blunted the reinforcement of somatosensory working memory (Sidarta et al., 2018), explaining the limited learning and consolidation observed in the subjects of Group_Long_. Another, potentially complementary interpretation is that rewards delivered after a long delay are temporally discounted and perceived as subjectively less valuable relative to when the delay is short (Shadmehr et al., 2019, 2010). If this is true, the reduction in subjective value in Group_Long_ may have contributed to reducing the beneficial effect of reward on offline memory consolidation (Sterpenich et al., 2021). Altogether, these elements indicate that the temporal contingency between a movement and the ensuing reward is a decisive aspect of reward-based motor learning.

Beyond reward timing, another feature that could have altered motor learning in the present study is the post-reward delay – *i*.*e*., the delay between reward delivery and the execution of the subsequent movement (referred to as ITI in the *Results* section, above). However, our data suggest that post-reward delay did not affect motor learning in the present study. Indeed, subjects of Group_Short_ and of Group_Short-PastStudy_, which trained with a short reward delay but with different ITIs, displayed very similar learning (Supplementary Figure 1). This finding is in line with the idea that motor learning remains largely unaffected by the ITI duration (even for ITIs of 30 s; (McDougle et al., 2015)).

In conclusion, our data indicate that the timing at which reward is delivered during motor training alters the dynamics of learning and the consolidation of the new motor memory. Research is now required to gain further knowledge as to the brain networks involved in these time-dependent effects of reward on motor learning as well as to characterize the relative contribution of implicit and explicit learning processes to these effects. Such knowledge would prove useful for the design of future reward-based rehabilitation programs, in which reward timing may be individualized depending on the brain networks and learning processes affected in specific populations of patients (Foerde et al., 2013; Foerde and Shohamy, 2011).

## 4. STAR METHODS

### RESOURCE AVAILABILITY

#### Lead contact

Further information and requests should be directed to the lead contact, Pierre Vassiliadis (contact).

#### Materials availability

This study did not generate new unique reagents.

#### Data and code availability

Data used in this study are available upon request (contact) and will be freely available at the time of publication via an open-access data sharing repository (https://osf.io).

### EXPERIMENTAL MODEL AND SUBJECT DETAILS

A total of sixty right-handed healthy volunteers participated in the present study (46 women, 23.7 ± 0.3 years old; mean ± SE). Data from a previous group of thirty participants was also re-analyzed (20 women, 23.9 ± 0.43 years old; (Vassiliadis et al., 2021)). Handedness was determined via a shortened version of the Edinburgh Handedness inventory (Oldfield, 1971). None of the participants suffered from any neurological or psychiatric disorder, nor were they taking any centrally-acting medication. All participants gave their written informed consent in accordance with the Ethics Committee of the Université Catholique de Louvain (approval number: 2018/22MAI/219) and the principles of the Declaration of Helsinki. Subjects were financially compensated for their participation. Finally, all participants were asked to fill out a French adaptation of the Sensitivity to Punishment and Sensitivity to Reward Questionnaire (SPSRQ; (Lardi et al., 2008; Torrubia et al., 2001)) and a NASA Task Load Index questionnaire (NASA-TLX, (Hart and Staveland, 1988)).

### METHOD DETAILS

#### Motor skill learning task

Participants were seated approximately 60 cm in front of a cathode-ray tube screen (refresh rate: 100 Hz) with their right forearm positioned at a right angle on the table. The task was developed on Matlab 7.5 (the Mathworks, Natick, Massachusetts, USA) exploiting the Psychophysics Toolbox extensions (Brainard, 1997; Pelli, 1997) and consisted in a previously described force modulation task (Vassiliadis et al., 2021). Briefly, the task required participants to squeeze a force transducer (Arsalis, Belgium) between the index and the thumb to control a cursor displayed on the screen. Increasing the force exerted resulted in the cursor moving vertically and upward. Each trial started with a preparatory period in which a sidebar appeared at the bottom of the screen and a target at the top (**Figure 1A**). After a variable time interval (0.8 to 1 s), a cursor popped up in the sidebar, indicating the start of the movement period. Participants had to pinch the transducer to move the cursor as quickly as possible from the sidebar to the target and maintain it there for the rest of the movement period, which lasted 2 s. The level of force required to reach the target (Target_Force_) was individualized for each participant and set at 10 % of maximum voluntary contraction (MVC). Notably, squeezing the transducer before the appearance of the cursor was considered as an anticipation and therefore led to the interruption of the trial. Anticipation trials were discarded from further analyses. At the end of each trial, a binary reinforcement feedback was presented to the subject (yellow or blue circle for success or failure, respectively).

#### Sensory and reinforcement feedbacks

We provided only limited visual feedback to the participants in order to increase the impact of the reinforcement feedback on learning (Mawase et al., 2017). As such, on 90 % of the trials, the cursor disappeared shortly after the start of the movement period: it became invisible as soon as the generated force became larger than half of the Target_Force_ (*i*.*e*., 5 % of MVC). Conversely, the remaining trials (10 % of the trials) provided a continuous vision of the cursor (full vision trials). Full vision trials were not considered in the analyses.

As mentioned above, each trial ended with the presentation of a binary reinforcement feedback, indicating success or failure. Success on the task was determined based on the Error, defined as the absolute force difference between the Target_Force_ and the exerted force (Abe et al., 2011; Steel et al., 2016). The Error was first computed for each frame refresh from 0.15 s to the end of the trial (*i*.*e*., providing 185 data points at 100 Hz), then averaged across the data points for each trial (Steel et al., 2016), and expressed in percentage of MVC. This indicator of performance allowed us to classify a trial as successful or not based on an individualized success threshold (see below). When the Error on a given trial was below the threshold, the trial was classified as successful, and when it was above the threshold, the trial was considered as failed. Hence, task success depended on the ability to approximate the Target_Force_ as quickly and as accurately as possible.

#### Reward timing manipulation

The protocol involved Training and Test blocks (see *Experimental protocol*, below). During Training blocks, reinforcement feedbacks were associated with a reward of 8 cents on successful trials, and failed trials led to 0 cent. Importantly, in two block types, we manipulated the timing at which the reinforcement feedback, and therefore the associated reward, was delivered after the movement period (**Figure 1A**). Indeed, the reward was displayed after either a short or a long delay – that is, 1 or 6 s following the movement period in Reward_Short_ and Reward_Long_ blocks, respectively (see (Foerde et al., 2013; Foerde and Shohamy, 2011) for the use of similar delays in decision-making tasks). In order to keep the total duration of the trial constant in these two block types, inter-trial intervals (ITI, which followed reward occurrence) were set to 6 and 1 s in the Reward_Short_ and the Reward_Long_ blocks, respectively. Finally, we re-analyzed data from a previous study (Vassiliadis et al., 2021), in which the training blocks involved a short reward delay timing (0.5 s) and an intermediate ITI (3 s; Reward_Short-PastStudy_ blocks). The latter analysis allowed us to test for the reproducibility of the effects of training obtained in the Reward_Short_ block.

In the Test blocks, reinforcement feedback occurred 1 s after the movement period, involved an ITI of 1 s, and was not associated with any reward.

#### Motor skill learning protocol

Subjects were tested on two consecutive days (Day 1 and Day 2; **Figure 1C**). On Day 1, we first measured the individual MVC to calculate the Target_Force_. Notably, MVCs and simple reaction times (SRT) were measured before and after the training blocks to assess potential fatigue related to the training (see **QUANTIFICATION AND STATISTICAL ANALYSIS**). Participants then performed 2 blocks of Familiarization, in a Test block setting. The first Familiarization block comprised 20 full vision trials. Subsequently, all blocks were composed of a mixture of partial vision trials (90 % of total trials) and full vision trials (10 % of total trials), as described above. The second Familiarization block involved 40 trials and allowed us to determine baseline performance to calibrate the difficulty of the task for the rest of the experiment (Calibration block; please see (Vassiliadis et al., 2021) for details on the Calibration procedure).

Following Familiarization, participants performed 320 trials divided in 8 blocks. All subjects started and ended the session with the realization of a Test block of 40 trials, allowing us to evaluate initial performance and total learning (*i*.*e*., Pre- and Post-training blocks, respectively). In between, 6 Training blocks (T1 to T6) of 40 trials were performed by the participants (see **Figure 1B**). During the Training blocks, individuals were split into 2 separate groups depending on the type of training blocks they performed. As such, Group_Short_ and Group_Long_ trained with Reward_Short_ and Reward_Long_ blocks, respectively. The group trained with Reward_Short-PastStudy_ blocks was referred to as Group_Short-PastStudy_. Comparing performance between the groups during the training period allowed us to test the effect of reward timing on the learning dynamics.

Day 2 was realized 24 hours later. Subjects performed the task again with the same Target_Force_ and success threshold. This assessment was composed of 5 full vision trials followed by a Test block of 40 trials (Re-test) and allowed us to assess the effect of reward timing on skill consolidation.

### QUANTIFICATION AND STATISTICAL ANALYSIS

Statistical analyses were carried out with Matlab 2018a (the Mathworks, Natick, Massachusetts, USA) and Statistica 10 (StatSoft Inc., Tulsa, Oklahoma, USA). In the case of independent samples t-tests and Analysis of Variance (ANOVA), we verified the homogeneity of the variances systematically and non-parametric tests were used when variances were non-homogeneous. All data were systematically tested for the sphericity assumption using Maunchley’s tests. The Greenhouse-Geisser (GG) correction was used for sphericity when necessary. Post-hoc comparisons were conducted using the Fisher’s LSD procedure. The significance level was set at p ≤ 0.05, except in the case of correction for multiple comparisons (see below).

#### Motor skill learning

As a first step, we tested the impact of reward timing on motor performance during each block of Test and Training block. We quantified for each subject the percentage of successful trials (*i*.*e*., the success rate) for each block and then normalized the data according to individuals’ initial performance by subtracting the success rate values measured at Pre-training from the values obtained in every block. To evaluate the impact of reward timing on success rates across training, we performed a two-way ANOVA with GROUP_TYPE_ (Group_Short_ and Group_Long_) and TRAINING_BLOCK_ (T1 to T6) as between- and within-subjects factors, respectively. Then, to evaluate the effect of the ITI’s duration on motor learning, we replicated this analysis with the inclusion of the Group_Short-PastStudy_.

As a second step, we aimed at evaluating how reward timing influenced learning rates during the task. To do so, we split the data into 24 non-overlapping bins of 10 trials, computed the success rate for each bin and normalized the data according to individuals’ initial performance, as done in the first analysis. The bins were then separated into two equal parts (*i*.*e*., of 12 bins each) depending on whether they belonged to the early or to the late phase of training (Training_Early_ and Training_Late_ phases, corresponding to T1-T3 and T4-T6, respectively). Finally, we performed linear regressions on these data and extracted the slope of the fits for the Training_Early_ and the Training_Late_ phases of the Group_Short_ and the Group_Long_. The slope values – exploited here as a proxy of the learning rate – were compared using a two-way ANOVA with GROUP_TYPE_ (Group_Short_ and Group_Long_) and TRAINING_PHASE_ (Training_Early_ and Training_Late_) as between- and within-subjects factors, respectively.

Finally, we tested for any effect of reward timing on total learning, by comparing the success rates of the three groups at Post-training, using a one-way ANOVA with the factor GROUP_TYPE_ (Group_Short_, Group_Long_ and Group_Short-PastStudy_). Further, in order to test the statistical significance of total learning within each group, we conducted two single sample t-tests on Post-training success rate, against a constant value of 0 (Bonferroni-corrected at p ≤ 0.025).

#### Motor skill consolidation

A secondary goal of the study was to evaluate the effect of reward timing on skill consolidation. For this analysis, we thus considered participants who demonstrated skill learning on Day 1 (Success_Post-training_ – Success_Pre-training_ > 0). This step not only allowed us to exclude non-learners but also to compare consolidation in groups that exhibited a similar learning on Day 1 (*i*.*e*., a similar success rate at Post-training). 40 participants were considered in this analysis (22 and 18 in Group_Short_ and Group_Long_, respectively). We compared the success rate data through a two-way ANOVA with GROUP_TYPE_ (Group_Short_ and Group_Long_) and TEST_BLOCK_ (Post-training and Day 2) as between- and within-subjects factors, respectively.

#### Group features, initial performance and fatigue

As a control, we verified that the Group_Short_ and the Group_Long_ were comparable in terms of age, success threshold, Target_Force_, sensitivity to reward and to punishment (*i*.*e*., as assessed by the SPSRQ questionnaire), initial performance (*i*.*e*., at Pre-training) and received monetary gains. As displayed in **Table 1**, independent sample two-tailed t-tests performed on these data did not reveal any significant differences between the groups (see also **Figure 1C**).

We also assessed if potential motor and cognitive fatigue generated by Day 1 training was different between the groups (Derosière et al., 2014; Derosiere and Perrey, 2012). To do so, we expressed MVCs, and SRTs obtained after training (MVC_POST_ and SRT_POST_) in percentage of the values measured initially (MVC_PRE_ and SRT_PRE_). We also assessed the perceived workload after training through the NASA-TLX questionnaire. Notably, these data did not differ between the groups (**Table 1**), suggesting that motor and cognitive fatigue were not responsible for the effect of reward timing on motor learning.

## Acknowledgements

P.V. was a PhD student supported by the Fund for Research training in Industry and Agriculture (FRIA/FNRS; FC29690), and grants by the Platform for Education and Talent (Gustave Boël - Sofina Fellowships) and Wallonie-Bruxelles International. J.D. was supported by grants from the Belgian FNRS and the Fondation Médicale Reine Elisabeth (FMRE). G.D. was supported by the Belgian FNRS and by ERC StG 2018 801872 Rhythm and Brains.

## Competing interests

The authors declare no conflict of interest.

**Supplementary Figure 1.**
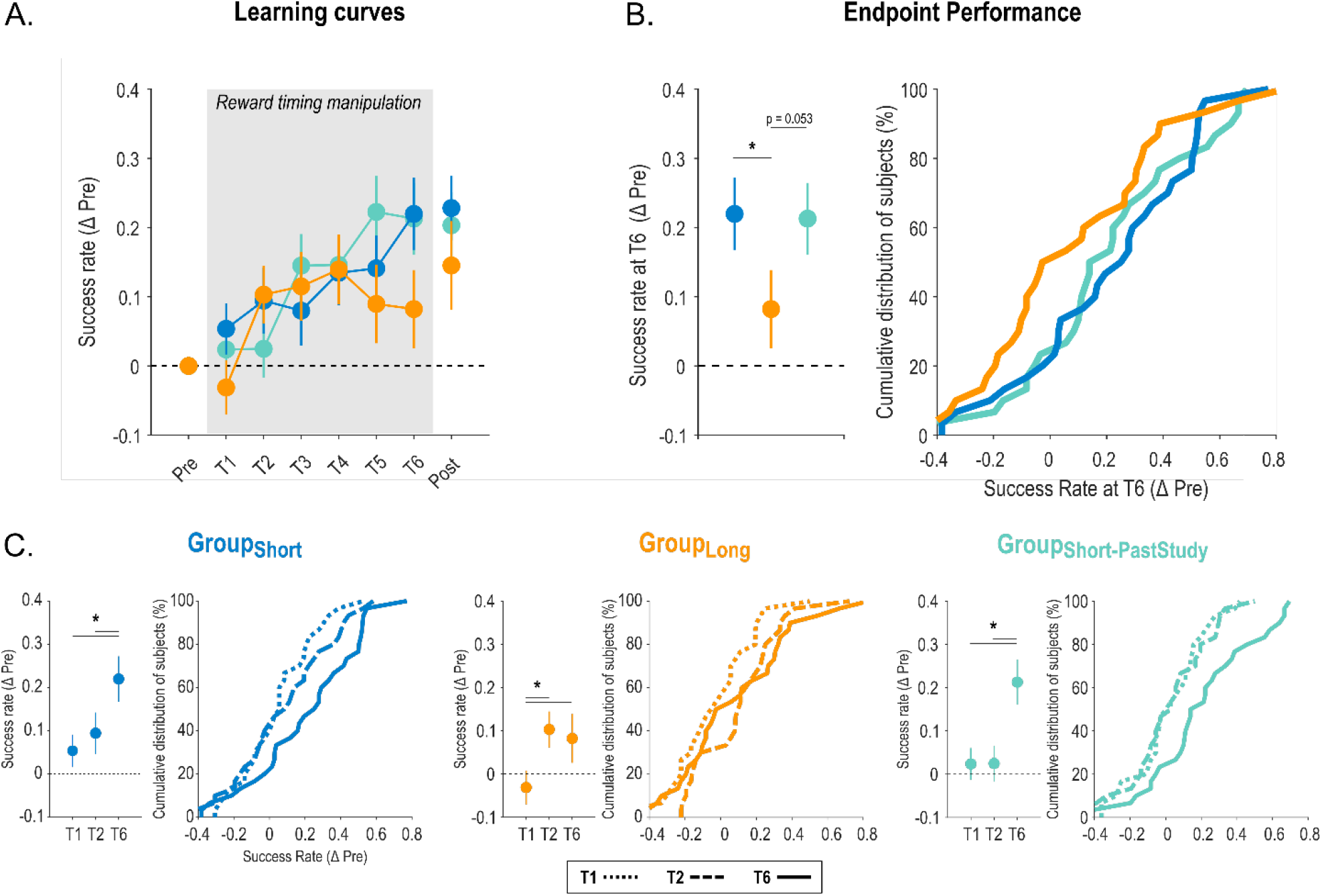
Effect of reward timing on motor skill learning with the inclusion of Group_PastStudy_. **A. Learning curves**. Proportion of successful trials (expressed as a difference with the individual Pre-training success rate) is represented across practice for the three experimental groups (blue: Group_Short_, orange: Group_Long_, turquoise: Group_Short-PastStudy_). The grey shaded area highlights the blocks concerned by the reward timing manipulation. The remaining blocks were Test blocks. Note the similarity in the learning curves of Group_Short_ and Group_Short-PastStudy_ despite the difference in ITI. **B. Endpoint performance**. Average success rates in the end of the training period (i.e., measured at T6) in the three groups (left panel) and the corresponding cumulative distributions of the data (right panel). **C. Learning dynamics**. Average success rates observed at T1 (dotted line), T2 (dashed line) and T6 (solid line) and the corresponding cumulative distributions for Group_Short_ (left), Group_Long_ (middle) and Group_Short-PastStudy_ (right panel). Performance improved very early in Group_Long_ (i.e., between T1 and T2) and then plateaus while it improved later in Group_Short_ and Group_Short-PastStudy_. Note that performance at T1 was not different between the groups (all p > 0.21). *: significant difference (p<0.05). Data are represented as mean ± SE.

